# Enhanced Synthesis of Poly Gamma Glutamic Acid by Increasing the Intracellular Reactive Oxygen Species in the *Bacillus licheniformis* Δ1-pyrroline-5-carboxylate Dehydrogenase Gene *ycgN* Deficient Strain

**DOI:** 10.1101/337402

**Authors:** Bichan Li, Dongbo Cai, Shiying Hu, Anting Zhu, Zhili He, Shouwen Chen

## Abstract

Poly gamma glutamic acid (*γ*-PGA) is an anionic polyamide with numerous applications. Proline metabolism influences the formation of reactive oxygen species (ROS), and is involved in a wide range of cellular processes. However, the relation between proline metabolism and *γ*-PGA synthesis has not yet been analyzed. In this study, our results indicated that the deletion of Δ1-pyrroline-5-carboxylate dehydrogenase encoded gene *ycgN* resulted in 85.22% higher yield of *γ*-PGA in *B. licheniformis* WX-02. But the deletion of proline dehydrogenase encoded gene *ycgM* had no effect on *γ*-PGA synthesis. Meanwhile, a 2.92-fold higher level of P5C was detected in *ycgN* deficient strain WXΔ*ycgN*, while the P5C levels in WXΔ*ycgM* and double mutant strain WXΔ*ycgMN* remained the same, compared to WX-02. The ROS level of WXΔ*ycgN* was 1.18-fold higher than that of WX-02, and the addition of n-acetylcysteine (antioxidant) into medium could decrease its ROS level, further reduced the *γ*-PGA yield. Our results showed that proline catabolism played an important role in maintaining ROS homeostasis, and the deletion of *ycgN* caused P5C accumulation, which induced a transient ROS signal to promote *γ*-PGA synthesis in *B. licheniformis*.

**Importance:** *γ*-PGA is an anionic polyamide with various applications in biomedical and industrial fields. Proline metabolism influences the intracellular reactive oxygen species (ROS) and is involved in a wide range of cellular processes. Here, we report the effects of proline metabolism on *γ*-PGA synthesis. Our results indicated that deletion of *ycgN* promoted the synthesis of *γ*-PGA by increasing the intracellular levels of Δ1-pyrroline-5-carboxylate to generate a transient ROS signal in *B. licheniformis* WX-02. This study provides the valuable information that enhanced synthesis of *γ*-PGA by knocking out of *ycgN*.

## Introduction

Poly gamma glutamic acid (*γ*-PGA) is an anionic polyamide which consists of D-and L-glutamic acid units connected by *γ*-amide linkages between *γ*-carboxyl and α-amino groups (1-3). With its features of hygroscopicity, water-solubility, biodegradability, non-toxicity and cation chelating, *γ*-PGA has been widely used in the fields of medicine, food, cosmetics, agriculture, water treatment, etc. (4). For example, *γ*-PGA can be served as the drug carrier in medicine, thickener and cryoprotectant in food industry, humectant in cosmetics, fertilizer synergist in agriculture, flocculants and heavy metal absorbent in water treatment (1, 4, 5).

*γ*-PGA is mainly produced by *Bacillus* species as an extracellular polymer (4, 6-8). Several strategies have been conducted to improve *γ*-PGA production via metabolic engineering of *γ*-PGA synthesis-related metabolic network (9). For instance, *γ*-PGA yield was enhanced by 63.2% in *B. amyloliquefaciens* LL3 via double-deletion of genes *cwlO* (encodes a cell wall lytic enzyme) and *epsA-O* cluster (responsible for extracellular polysaccharide synthesis), as well as introduction of the gene *vgb* (encodes *Vitreoscilla* hemoglobin) (10). Another example is that a systematically metabolic engineering study consisting the by-products synthesis, *γ*-PGA degradation, glutamate precursor synthesis, *γ*-PGA synthesis and autoinducer synthesis pathways has been performed in *B. amyloliquefaciens* NK-1, and the *γ*-PGA yield was increased by 2.91-fold in the final strain NK-anti-*rocG* (11). In our previous work, over-expression of *glr* (encodes glutamic acid racemase) (12), *pgdS* (encodes *γ*-PGA hydrolase) (3), *zwf* gene (encodes glucose-6-phosphate dehydrogenase) (13), *rocG* (encodes glutamate dehydrogenase) (14), and *fnr* (encodes global anaerobic regulator) (15) could all enhance *γ*-PGA production in *B. licheniformis* WX-02. Also, improving the capability of assimilating glycerol was proved to be an efficient strategy to increase *γ*-PGA yield, and the *γ*-PGA concentration was improved by 33.71% by substituting the native *glpFK* promoter with the constitutive promoter P43 (16).

Proline is a multifaceted amino acid with important roles in carbon and nitrogen metabolism, protein synthesis, bioenergetics, differentiation, growth, etc. (17-19). Proline is oxidated to glutamate by proline dehydrogenase (PRODH) and Δ1-pyrroline-5-carboxylate dehydrogenase (P5CDH), involved a four-electron oxidation process (**Fig. 1**) (18). Firstly, proline is oxidized to Δ1-pyrroline-5-carboxylate (P5C) by O_2_-dependent PRODH, which is non-enzymatically transformed into glutamate-*γ*-semialdehyde (GSA), and then oxidized to glutamate by P5CDH (18, 20). Recent studies demonstrated that proline metabolism plays an important role in maintaining the balance of intracellular reactive oxygen species (ROS), and influenced numerous additional regulatory pathways, such as p53-mediated apoptosis in mammalian cells (21) and osmo-protecting mechanisms in plants(17) and bacteria(18, 22), etc. There are two possible mechanisms for the ROS generation. First, ROS is supposed to be generated from P5C-proline cycle, which is catalyzed by PRODH and P5C reductase (**Fig1**) (17, 23, 24). The P5C-proline cycle provides an excess of electron flow to the electron transport chain (ETC) and O_2_ which further induces ROS overproduction (17, 23, 24). Second, ROS is spontaneously generated from the intermediate of proline metabolism P5C/GSA, which has the high activity with various cellular compounds (25, 26). In budding yeast, P5C directly inhibits the mitochondrial respiration and induces a burst of superoxide anions from the mitochondria (24). Since addition of H_2_O_2_ was proven as an efficient strategy for enhancement of *γ*-PGA yield by improving the intracellular ROS (9), we hypothesized that the manipulating of proline metabolism might also affect *γ*-PGA synthesis.

**Fig. 1.**
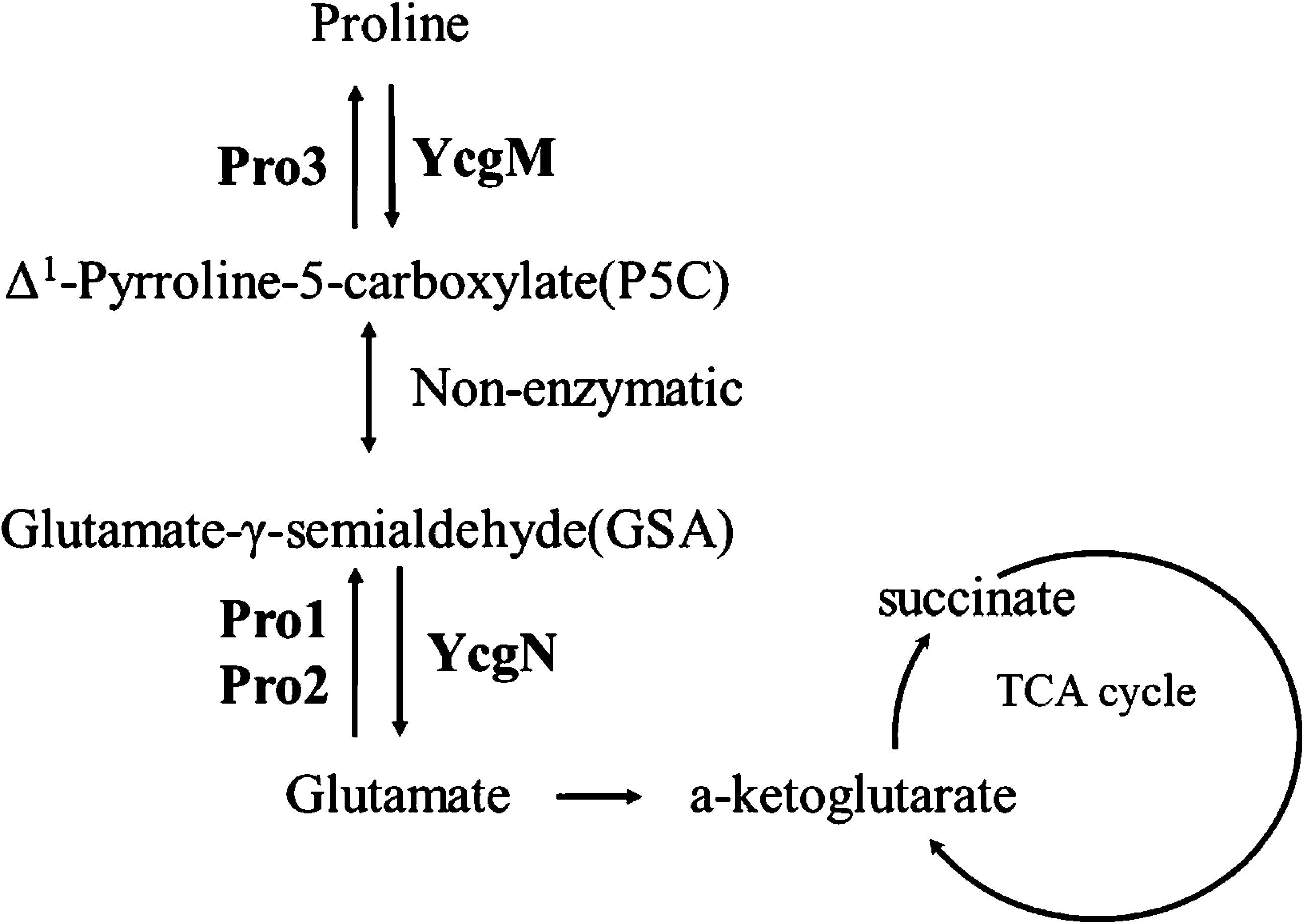
The scheme of proline degradation pathway in *B. licheniformis* WX-02. Pro1, *γ*-glutamyl kinase; Pro2, *γ*-glutamyl phosphate reductase; Pro3, P5C reductase; YcgM, proline oxidase; ycgN, P5C dehydrogenase; TCA cycle, the tricarboxylic acid cycle.

Reactive oxygen species (ROS), such as superoxide 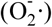, hydrogen peroxide (H_2_O_2_), and hydroxyl radical (OH·), are highly reactive molecules. They mediate a number of significant cell processes, including impaired cellular homeostasis, enzyme inactivation, DNA damage and cell death (27). ROS could also implicate in pathologies, such as cancer, atherosclerosis, diabetes, Down’s syndrome, and in neurodegenerative diseases like Parkinson’s and Alzheimer’s diseases (25, 27, 28). In addition, ROS could serve as the secondary messengers to activate the downstream defense against invaded microorganisms in plants (29, 30), and played an important role in plant secondary metabolism (31). Moreover, ROS could regulate the product synthesis in microorganism, and the yields of xanthan gum produced by *Xanthomonas campestris* (29), validamycin A produced by *Streptomyces hygroscopicus* 5008 (31), *γ*-PGA produced by *Bacillus subtilis* NX-2 (9) and fumonisin produced by *Fusarium verticillioides* (32, 33) were all improved obviously by the ROS induction.

*B. licheniformis* WX-02 has been proven to be an efficient *γ*-PGA producer (13), and several strategies have been conducted to improve *γ*-PGA synthesis. The yield of *γ*-PGA was increased 2.3-fold by addition of nitrate in the medium (2). Physicochemical stresses such as heat, osmotic and alkaline could enhance the production of *γ*-PGA (34, 35). In this study, *ycgM* (encodes PRODH) and *ycgN* (encodes P5CDH) were deleted respectively to analyze the role of proline metabolism on *γ*-PGA synthesis. The intracellular concentrations of proline, P5C and ATP, ROS levels were analyzed, and the related gene transcriptional levels were measured, to expound the mechanism of *ycgN* deletion on *γ*-PGA synthesis.

## RESULTS

### Deletion of *ycgN* improved *γ*-PGA production in *B. licheniformis*

The catabolism of proline was catalyzed by YcgM (PRODH) and YcgN (P5CDH) in *B. licheniformis* WX-02(**Fig. 1**). To investigate the effects of proline metabolism on *γ*-PGA yield, *ycgM* and *ycgN* were deleted in WX-02, and resulted in strains WXΔ*ycgM* and WXΔ*ycgN*, respectively. These recombinant strains, as well as the control strain WX-02, were cultivated in the *γ*-PGA production medium. As shown in **Fig. 2A**, the *γ*-PGA yield of WXΔ*ycgM* was 7.58 g L^-1^. The *γ*-PGA yield of WXΔ*ycgN* was 13.91g L^-1^, which was 85.22% higher than that of WX-02 (7.51 g L^-1^). The yield of *γ*-PGA from the complementation strain WXΔ*ycgN-*N had no difference with WX-02 (**Fig. 2B**), which confirmed the depletion of *ycgN* along contributes to the increase of *γ*-PGA yield. To dissect the effect of *ycgN* deletion on *γ*-PGA synthesis, the *ycgM* and *ycgN* double mutant strain WXΔ*ycgMN* was constructed, which was expected to lack the potential toxic compounds P5C/GSA (**Fig. 1**). Based on our results, the *γ*-PGA yield of WXΔ*ycgMN* was 7.59 g L^-1^, similar to that of WX-02 (**Fig. 2B**). Furthermore, the *ycgM* and *ycgN* genes were overexpressed in WX-02, resulting in WX/*ycgM* and WX/*ycgN*, respectively. As shown in **Fig. 2B,** the *γ*-PGA yields of WX/*ycgM* (8.49 g L^-1^) and WX/*ycgN* (8.39 g L^-1^) were lower than that of control strain WX/pHY300 (11.82 g L^-1^).

**Fig. 2.**
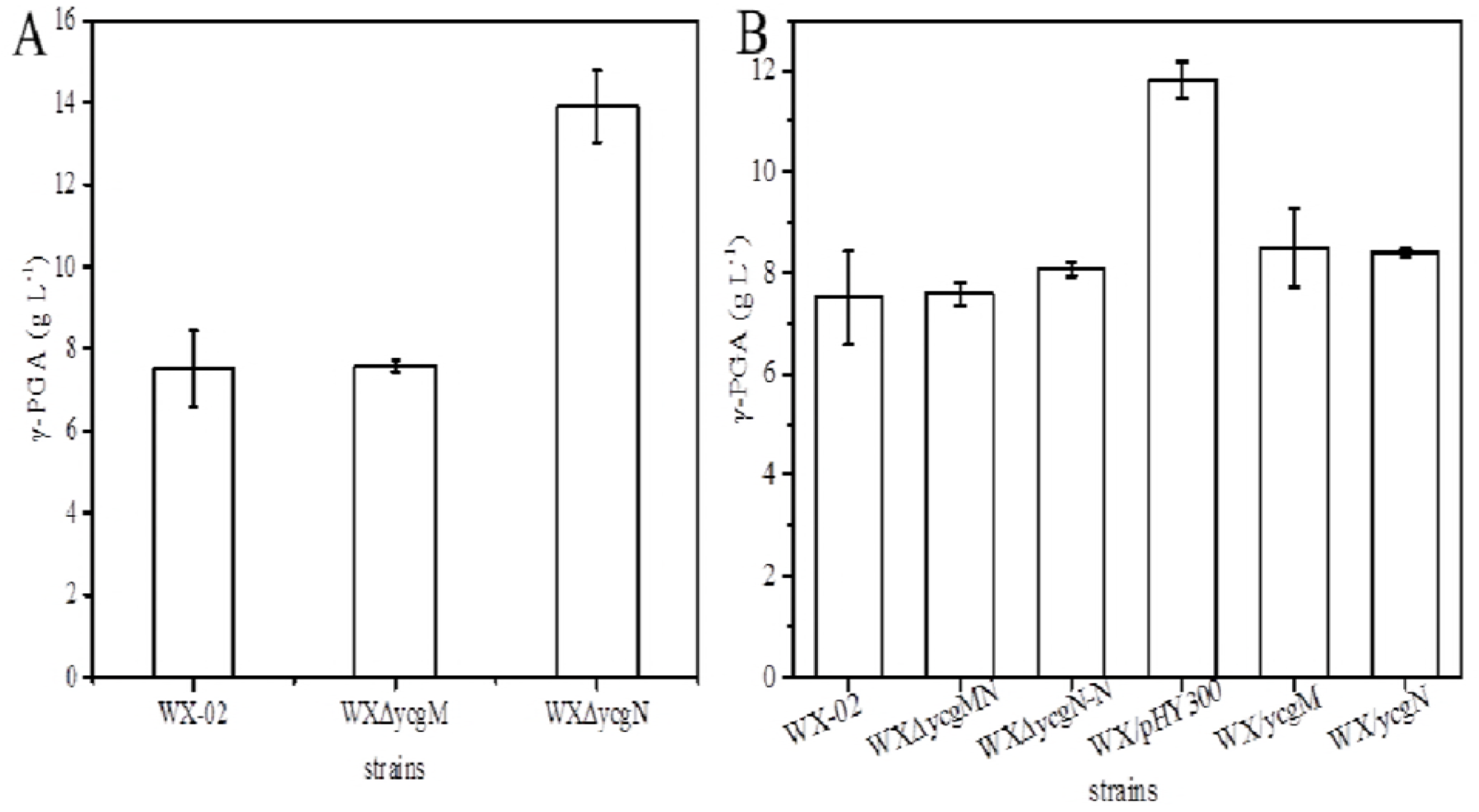
The *γ*-PGA yields of *ycgM* and *ycgN* deletion strains (A); complements strains WX△*ycgN*-N, WX△*ycgMN*, WX/*ycgM* and WX/*ycgN* (B).

The *γ*-PGA yields, biomass, and glucose concentrations during the *γ*-PGA synthesis were investigated to observe the influences of deletions of *ycgM*, *ycgN*, and double deletion of *ycgMN*. As shown in **Fig. 3,** no remarkable changes of *γ*-PGA yield, cell growth, or glucose consumption rate were observed among WXΔ*ycgM*, WXΔ*ycgMN* and WX-02. However, the yield of *γ*-PGA from WXΔ*ycgN* was increased by 85.22% (**Fig. 3A**). The lag phase of WXΔ*ycgN* was 4∽8 hours longer, and the exponential phase of WXΔ*ycgN* was 4 hours longer. The maximal biomass of WXΔ*ycgN* was 14.07% higher than that of WX-02(**Fig. 3B**). The glucose consumption rate of WXΔ*ycgN* was 1.52 g L^-1^ h^-1^, which was 22.44% lower than that of WX-02(1.96 g L^-1^ h^-1^) (**Fig. 3C**).

**Fig. 3.**
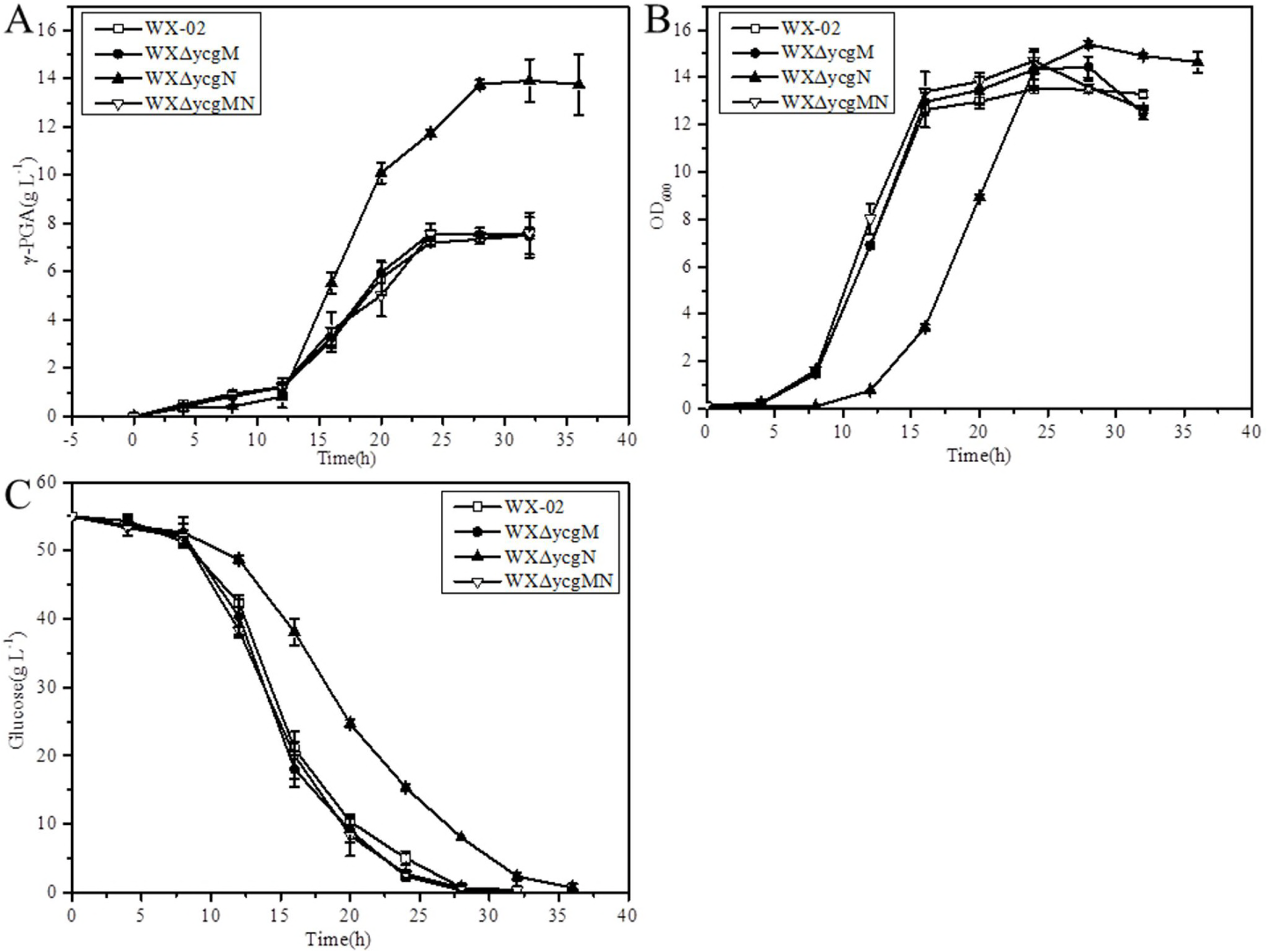
Comparison of *γ*-PGA synthesie (A), cell growth (B) and glucose consumption (C) of *ycgM* and *ycgN* deletion strains during *γ*-PGA production. Values are averages from three biological replicates.

### Effect of *ycgN* deficiency on the intracellular proline and P5C concentrations

In *B. licheniformis*, proline was degraded to P5C by YcgM. P5C was then spontaneously changed to GSA, and further converted to glutamate by YcgN (**Fig. 1**). Higher proline levels were observed in both WXΔ*ycgM* (10.04 μmol·gDCW^-1^) and WXΔ*ycgMN* (8.95 μmol·gDCW^-1^) (**Fig. 4**). Whereas, the proline concentration of WXΔ*ycgN* (6.68 μmol·gDCW^-1^) was similar to that of WX-02 (6.26 μmol·gDCW^-1^). Meanwhile, the P5C accumulated in WXΔ*ycgN* (19.24 μmol·gDCW^-1^) was significantly higher than that of WX-02 (4.91 μmol·gDCW^-1^) (**Fig. 4**). And the P5C concentration of WXΔ*ycgM* (4.76 μmol·gDCW^-1^) and WXΔ*ycgMN* (4.92 μmol·gDCW^-1^) were comparable to that of WX-02 (**Fig. 4**).

**Fig. 4.**
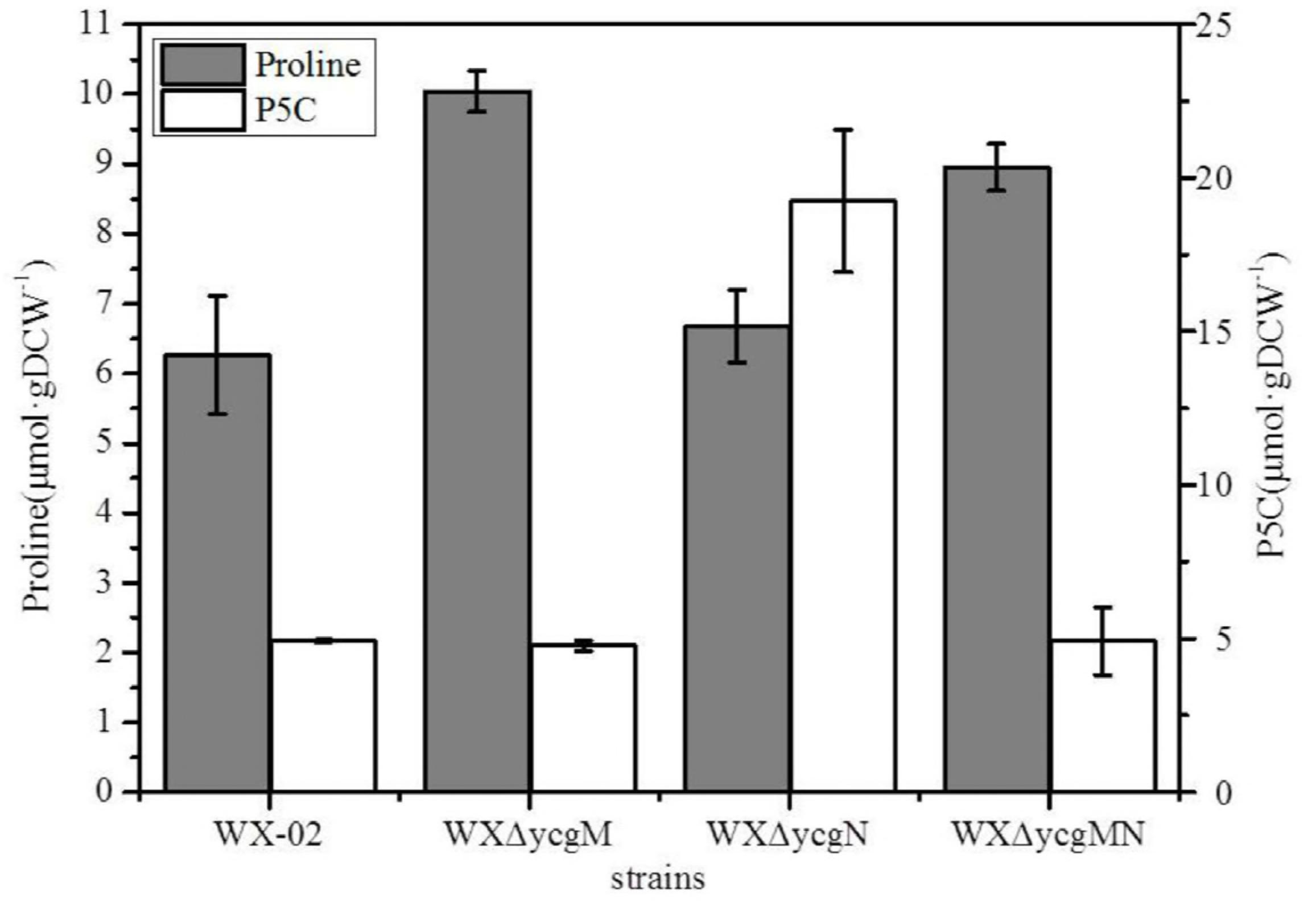
The intracellular concentrations of proline (A) and P5C (B) in *ycgM* and *ycgN* deletion strains. Data represent the mean and standard deviation from three independent experiments. DCW means dry cell weight.

### The intracellular ROS levels was enhanced in the *ycgN* deletion strain

P5C was reported to be a direct inhibitor of mitochondrial respiration in yeast, which might further lead to intracellular ROS accumulation (24). Thus, we hypothesized that the deletion of *ycgN* might enhance the accumulation of intracellular ROS, which further influenced *γ*-PGA synthesis. To confirm this hypothesis, the ROS levels of WX-02, WXΔ*ycgM*, WXΔ*ycgN*, and WXΔ*ycgMN* were measured by DCFH method. As shown in **Fig. 5A**, a 1.18-fold increase in fluorescence was detected in WXΔ*ycgN* at 4 h. And the ROS content in WXΔ*ycgN* was found to be significantly decreased after 8 h (**Fig. 5A**). The intracellular ROS levels in WXΔ*ycgM* and WXΔ*ycgMN* were 42.60% and 29.38% lower than that of WX-02, respectively (**Fig. 5A**). These results proposed that deletion of *ycgN* induced a transient increase in ROS, which might promote ***γ*-PGA** synthesis.

**Fig. 5.**
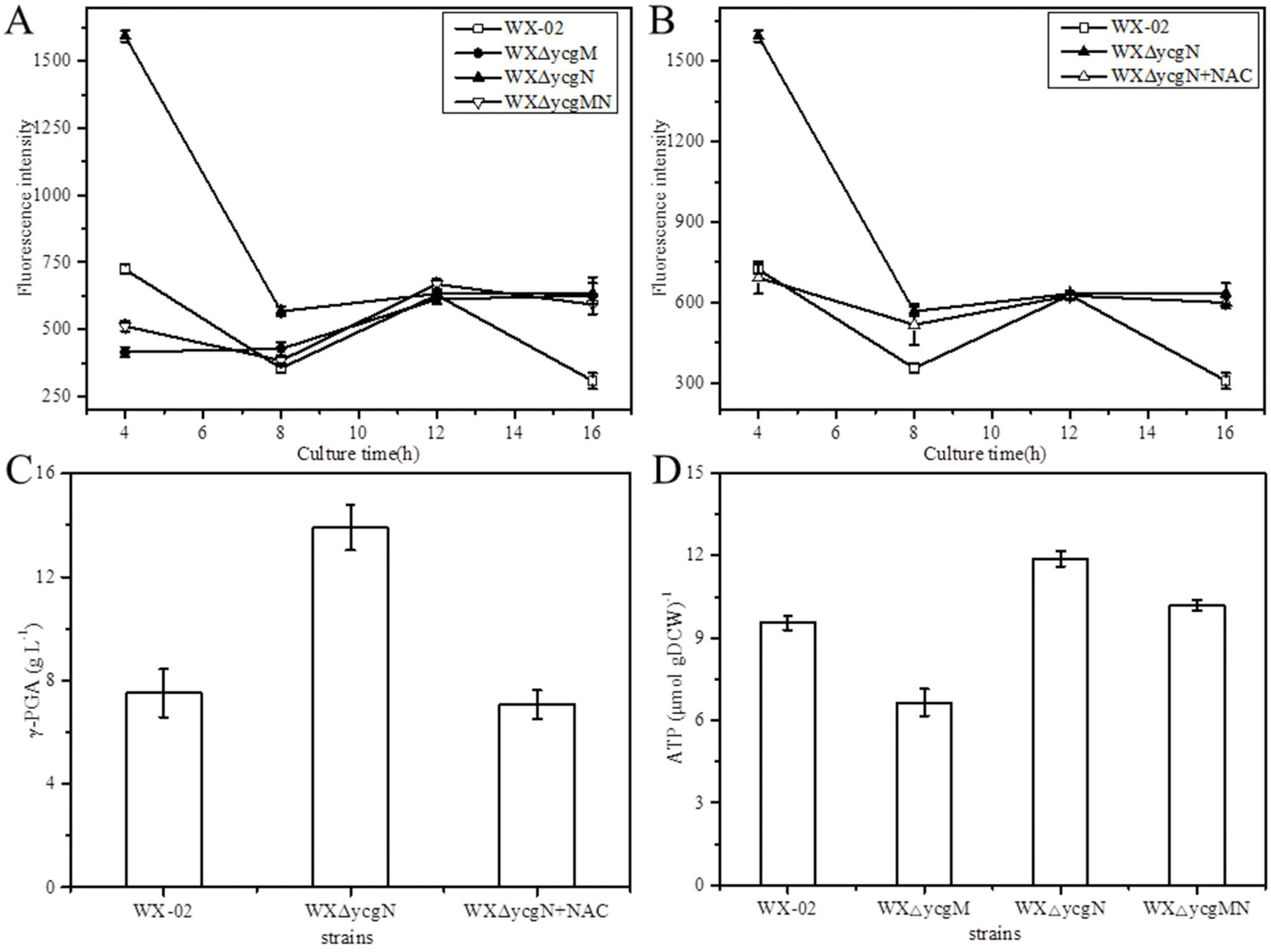
The intracellular concentrations of ROS in the mutants and its effects on *γ*-PGA synthesis. The intracellular concentrations of ROS in *ycgM*, *ycgN* single mutant and double mutants (A). Effects of exogenous antioxidant addition on the level of intracellular ROS (B). Effects of exogenous antioxidant addition on *γ*-PGA synthesis (C). The intracellular ATP concentrations of mutant strains during *γ*-PGA fermentation (D).

### The *ycgN*-dependent ROS signal contributed to *γ*-PGA synthesis

ROS has been reported to promote *γ*-PGA synthesis capability in *B. subtilis* NX-2 previously (9). Thus, the transiently increased ROS level observed in WXΔ*ycgN* was proposed to be the primary cause of *γ*-PGA enhancement. To test this hypothesis, n-acetylcysteine (NAC, antioxidant) (10 mmol L^-1^), a widely used antioxidant agent, was added into the medium to neutralize the ROS production. In the present of NAC, the ROS level of WXΔ*ycgN* was similar to that of WX-02 at 4 h (**Fig. 5B**). The *γ*-PGA yield of WXΔ*ycgN* was 7.06 g L^-1^, which is near to WX-02 (**Fig. 5C**). To further verify that ROS would promote *γ*-PGA synthesis in *B. licheniformis* WX-02, H_2_O_2_ was supplied as a simple mean to increase the intrinsic ROS levels. Our result implied that addition of 10 mmol L^-1^ H_2_O_2_ could increase the *γ*-PGA yield (13.88 g L^-1^) by 77.72% (**Fig. S1**). Collectively, our results suggested that the increase of ROS induced by the deletion of *ycgN* might contribute to the enhancement of *γ*-PGA yield in WXΔ*ycgN*. Overproduction of ROS has been reported to cause damage of intracellular biomolecules, such as proteins, DNA and lipids, which was not conducive to the cell growth (25, 39, 40). Consequently, WXΔ*ycgN* exhibited a slight growth defect and the extended lag phase, compared with those of WX-02.

### Effect of *ycgN* deletion on the intracellular ATP concentration

ATP supply is essential for product synthesis, as well as in *γ*-PGA (15). As show in **Fig. 5D**, the intracellular ATP concentration of WXΔ*ycgN* was 11.875 μmol·gDCW^-1^, increased by 24.40% compared to that of WX-02 (9.55 μmol·gDCW^-1^). The ATP concentrations of WXΔ*ycgM* and WXΔ*ycgMN* were 6.65 μmol·gDCW^-1^ and 10.19 μmol·gDCW^-1^, respectively. These results indicate that deletion of *ycgN* improved the intracellular ATP supply, which was beneficial for *γ*-PGA synthesis.

### Transcriptional levels of genes related to *γ*-PGA synthesis in *ycgN* mutant strain

The general stress response of *B. licheniformis* is controlled by the *σ*^B^ transcription factor encoded by *sigB* (41). To approve that deletion of *ycgN* could cause oxidative stress in *B. licheniformis* WX-02, the transcription level of *sigB* was determined in WXΔ*ycgN*. As shown in **Fig. 6**, the relative expression level of the gene *sigB* in WXΔ*ycgN* was increased to 13.15.

**Fig. 6.**
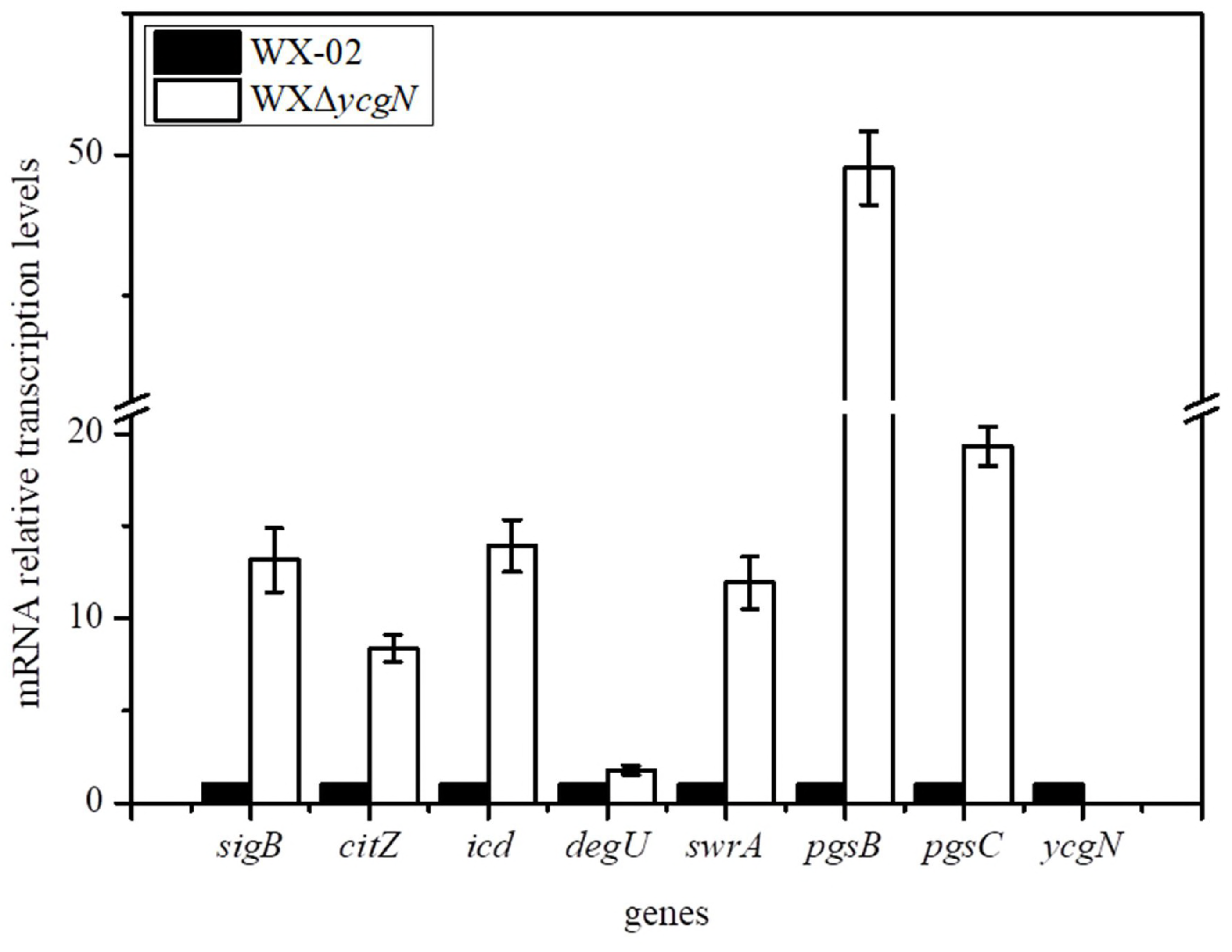
Effects of *ycgN* delection on the relative transcriptional levels of genes in TCA cycle and *γ*-PGA biosynthesis.

To investigate the roles of YcgN on expression of genes involved in *γ*-PGA synthesis, the transcription levels of *degU, swrA, pgsB*, and *pgsC* were analyzed in WXΔ*ycgN*. As shown in **Fig. 6**, the transcriptional levels of genes *pgsB* and *pgsC*, which are responsible for *γ*-PGA biosynthesis, were increased by 49.52-and 19.31-fold, respectively. The expression of *pgs* operon is activated by SwrA and phosphorylated DegU (DegU-P) (4). Accordingly, the transcriptional levels of genes *degU* and *swrA* were enhanced by 1.79-and 11.92-fold, respectively (**Fig. 6**). Also, the transcriptional levels of relevant genes in TCA cycle, including *citZ* (encodes citrate synthase) and *icd* (encodes isocitrate dehydrogenase) were verified, as the precursor of *γ*-PGA glutamate can be synthesized from α-Ketoglutaric acid. And the expression levels of both genes were increased by 8.36-and 13.93-fold in WXΔ*ycgN*, respectively (**Fig. 6**).

### Discussion

Proline is a multifunctional amino acid which can be used as carbon, nitrogen and energy source (18). Also, it plays an important role in protecting against osmotic and oxidative stresses, since it is a compatible solute and a free-radical scavenger (42). In addition, proline catabolism has been found to be involved in protection of intracellular redox homeostasis and virulence in microorganisms (26, 43). In this study, we demonstrated that the deletion of *ycgN* significantly enhanced *γ*-PGA production in *B. licheniformis* WX-02, and this phenomenon was disappeared in the complementation strain. Besides, the increases of P5C concentration, ROS and intracellular ATP concentration were observed in *ycgN* deletion strain. These results indicated that proline metabolism is valuable in regulating *γ*-PGA synthesis, and the transient increase in ROS level seemed to be required for the enhancement of *γ*-PGA synthesis caused by deletion of *ycgN*.

Briefly, the degradation of proline to glutamate is catalyzed by PRODH and P5CDH. Glutamate can be converted to α-ketoglutarate (α-KG) by a transamination, and then oxidized in the TCA cycle along with ATP generation (**Fig. 1**). In the previous research, knocking out of P5CDH encoded gene *Ldp5cdh* in the Colorado potato beetle *Leptinotarsa decemlineata* could significantly reduce the ATP content, and further inhibit flight capacity (42). This study implied that deletion of *ycgN* in *B. licheniformis* WX-02 led to an 85.22% increase of *γ*-PGA yield. One explanation is that deletion of *ycgN* prevents the pathway of proline oxidation, and then reduces ATP content. However, the catabolism of proline was interrupted in *ycgM* mutant and *ycgMN* double mutant strains (Fig. 1), the *γ*-PGA yield exhibited no difference from the wild-type. Moreover, the ATP content of *ycgN* deletion strain was significantly higher than that of wild-type strain and other mutants. Thus, the interrupting of proline oxidation and ATP generation might not be the main reason for the enhancement of *γ*-PGA yield obtained in the *ycgN* deletion strain.

Based on our results, the enhancement of *γ*-PGA in WXΔ*ycgN* was related to the ROS accumulation (**Fig. 5**). In the previous researches, impaired P5C dehydrogenase activity was supposed to induce ROS generation by causing intense P5C-proline cycling in animals, plants and fungus (26). The oxidation of proline to P5C by FAD dependent-ProDH provides an excess of electrons to the mitochondrial electron transport chain and enhances ROS accumulation (17, 26, 44). P5CDH would prevent the excessive producing and accumulation of ROS by converting P5C to glutamate irreversibly (44). However, overexpression of *ycgM* did not improve *γ*-PGA production in WX-02, indicating that the increase of *γ*-PGA production in WX-02Δ*ycgN* was not attributed to the proline-P5C cycling. Another interpretation was that the effects of *ycgN* mutation were mediated by P5C. Consistently, an increase of P5C was detected in *ycgN* mutant but not in other mutants. Hence, the improvement of *γ*-PGA production in *ycgN* deletion strain might due to the P5C or P5C-derived signals.

Formaldehyde and acetaldehyde, which containing the aldehyde group, had been proved to be able to impair mitochondrial function and then generate ROS (45, 46). P5C attacks the mitochondrial respiratory chain and induces a burst of superoxide anions from the mitochondria in *Saccharomyces cerevisiae* R1278b (24). P5C is also a primary inducer of p53-mediated apoptosis and ROS-dependent autophagy proposed in mammals (47). In plants, P5CDH infection resulted in P5C accumulation and induced ROS burst (48, 49). Thus, it might be reasonable that P5C or, more likely, its equilibrium compound GSA with an unstable aldehyde group, contributed to the *γ*-PGA enhancement by inducing ROS burst via inhibiting the respiratory chain. Based on our results, a transient increase of ROS was observed at earlier time points in WX-02Δ*ycgN* mutants, but not in WX-02, WX-02Δ*ycgM* or WX-02Δ*ycgMN*.

*γ*-PGA is a homopolymer of glutamate with diverse biochemical properties (4). Several organisms secrete *γ*-PGA into the environment for sequestration of toxic metal ions or decreasing high local salt concentrations, enabling them to survive in adverse conditions (4). In our previous researches, the *γ*-PGA synthesis capability was strengthened when the strains were cultured in the stress conditions, such as high salt, high temperature, caustic alkali, and ultrasonic shock (20). Here, it was found that the *γ*-PGA synthesis of WX-pHY300 was increased by 57.40% compared with that of WX-02 (**Fig. 2**), which was in line with the previous studies. Since the plasmid was supposed to exhibit metabolic burden on the host and affected host gene expression and phenotype, it was suspicious that the enhancement of *γ*-PGA synthesis in WX-pHY300 could be an element of response or adaptation response against stress caused by pHY300 (50-53). According to this study, oxidative stress increased *γ*-PGA production in WX-02 by *ycgN* deletion or H_2_O_2_ addition. The addition of n-acetylcysteine decreased the *γ*-PGA yield of WXΔ*ycgN* to the level near that of WX-02. Thus, *γ*-PGA synthesis could be an element for adaptation response against oxidative stress.

ROS has been proposed to act as the secondary messenger and regulate many processes at the transcriptional level (19, 31, 54, 55). The global regulator OxyR was reported to react with H_2_O_2_ and form a disulfide bond between Cys199 and Cys208, resulting in the transcriptional activation of OxyR regulator in *E. coli* (19). In *B. subtilis*, ROS, induced by high shear stress, altered the transcription of general protein Sigma B and Ctc, and then regulated the suppression of sporulation (31). The transcription levels of *degU, swrA*, and *pgsB* which are related to *γ*-PGA biosynthesis were markedly increased in the *ycgN* deletion strain. In recent researches, DegU was proposed to be under control of the redox-sensing regulators ClpXP/Spx (56-58). Therefore, the intracellular ROS, induced by *ycgN* deletion, promoted the transcription level of *degU* probably by activating Spx. Also, the transcription of *swrA* was significantly improved in WXΔ*ycgN*, and the improvement of SwrA might cooperate with DegU to active the expression of *pgs* operon and promote *γ*-PGA synthesis (**Fig. 7**).

**Fig. 7.**
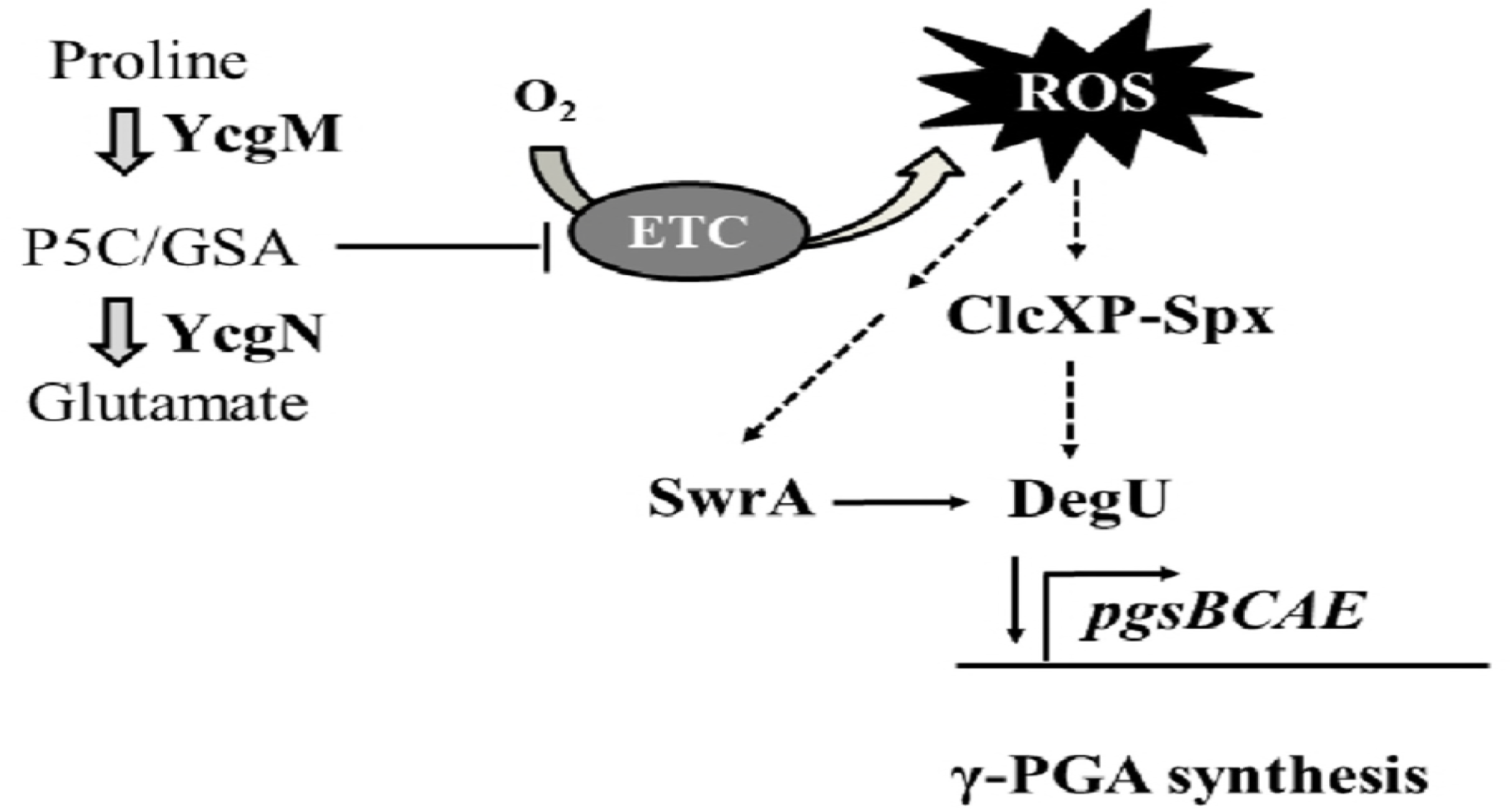
The proposed mechanism of the influence of proline metabolism on *γ*-PGA synthesis. Arrows indicate activation or promotion; T bars indicate repression or inhibition. Solid lines indicate empirical supports for regulation, and dashed lines indicate supports by inferences where the mechanism of regulation is unknown.

### Conclusion

The role of proline metabolism on *γ*-PGA synthesis is analyzed in this work. Based on our results, *γ*-PGA synthesis in *B. licheniformis* WX-02 was enhanced by the deletion of *ycgN*, which yield was 85.22% higher than that of wild-type strain. Secondly, the P5C concentration of WX-02Δ*ycgN* was 2.92-fold increased, which resulted in the intracellular ROS accumulation. These results illustrate the importance of P5C dehydrogenase in regulating *γ*-PGA production, and it provides valuable information for metabolic engineering of high-yield *γ*-PGA strain of *B. licheniformis*.

## MATERIALS AND METHODS

### Bacterial strains, media and culture conditions

The strains and plasmids used in this work are listed in **Table 1**. *B. licheniformis* and *Escherichia coli* were cultured at 37°C in Luria-Bertani (LB) broth (1% tryptone, 0.5% yeast extract, 1% NaCl and pH 7.2). The seed culture of *B. licheniformis* was prepared in a 250 mL flasks containing 50 mL LB medium, and incubated at 37°C in a rotatory shaker (180 rpm) for 10 h until OD_600_ reached 4.6∽5.0. The seed culture (1.50 mL) was inoculated into 250 mL flask containing 50 mL *γ*-PGA production medium (g L^-1^: glucose 60, sodium nitrate 10, sodium citrate 10, NH_4_Cl 8, CaCl_2_ 1, K_2_HPO_4_·3H _2_O 1, MgSO_4_·7H _2_O 1, ZnSO_4_·7H _2_O 1, MnSO_4_·7H _2_O 0.15 and FeCl_3_·6H _2_O 0.04), and shaken at 37°C and 180 rpm for 32 h. All the fermentation experiments were performed in three replicates. The antibiotics kanamycin and tetracycline were added into *B. licheniformis* cultures at the final concentration of 20 mg L^-1^ when necessary. Kanamycin and Ampicillin were added to *E. coli* cultures at final concentrations of 20 mg L^-1^ and 50 mg L^-1^, respectively.

**Table 1.** The strains and plasmids used in this study.

| Strains or plasmids | Description | Source |
| --- | --- | --- |
| **Strains** |  |  |
| ***B. licheniformis*** |  |  |
| WX-02 | Polyglutamate productive strain (CCTCC M208065) | Laboratory stock |
| WXΔycgM | *B. licheniformis* WX-02 carrying an in-frame deletion in ycgM gene | This study |
| WXΔycgN | *B. licheniformis* WX-02 carrying an in-frame deletion in ycgN gene | This study |
| WXΔycgMN | *B. licheniformis* WX-02 carrying an in-frame deletion in ycgM and ycgN gene | This study |
| WX/pHY300 | WX-02 harboring pHY300PLK | This study |
| WX/ycgM | WX-02 harboring pHY-ycgM | This study |
| WX/ycgN | WX-02 harboring pHY-ycgN | This study |
| WXΔycgN-N | WXΔycgN complemented with ycgN inserted in the genome at amyL locus. | This study |
| ***Escherichia coli*** |  |  |
| DH5α | F<sup>-</sup> Φ80d/lacZΔM15, $\Delta$(lacZYA-argF)U169, recA1, endA1, hsdR17(rK<sup>-</sup>, mK<sup>+</sup>), phoA, supE44, $\lambda^-$, thi-1, gyrA96, relA1 | Laboratory stock |
| **Plasmids** |  |  |
| T2(2)-ori | *E. coli*-*B. licheniformis* shuttle vector, OripUC/Orits, Kan | Laboratory stock |
| T2-ycgM | T2(2)-ori derivative, carrying homology arms for the deletion of ycgM gene | This study |
| T2-ycgN | T2(2)-ori derivative, carrying homology arms for the deletion of ycgN gene | This study |
| T2-ycgMN | T2(2)-ori derivative, carrying homology arms for the deletion of ycgM and ycgN gene | This study |
| T2-P<sub>srf</sub>-ycgN | T2(2)-ori derivative, carrying amyL:: (P<sub>srf</sub>-ycgN) | This study |
| pHY300PLK | *E. coli*-*B. licheniformis* shuttle vector, Ap<sup>r</sup>(*E. coli*), Tc<sup>r</sup>(*E. coli* and *B. licheniformis*) | This study |
| pHY-ycgM | pHY300PLK containing P<sub>srf</sub> promoter, the gene ycgM and amyL terminator | This study |
| pHY-ycgN | pHY300PLK containing P<sub>srf</sub> promoter, the gene ycgN and amyL terminator | This study |

### Construction of plasmids and strains

DNA manipulations were performed according to our previous researches (13, 36). The construction procedure of *ycgN* deficient strain was served as an example. Briefly, the homology arms of gene *ycgN* were amplified from chromosomal DNA of *B. licheniformis* WX-02 with primers *ycgN*-A – F/*ycgN*-A-R and *ycgN*-B – F/*ycgN*-B-R (**Table 2**), respectively. The resulting fragments were purified and ligated by Splicing Overlapping Extension PCR (SOE-PCR) with the primers *ycgN*-A-F and *ycgN*-B-R. The fused fragment was digested with *Bam*HI and *Xba*I, and inserted into T2(2)-Ori, named T2-*ycgN*(13, 36).

**Table 2.**
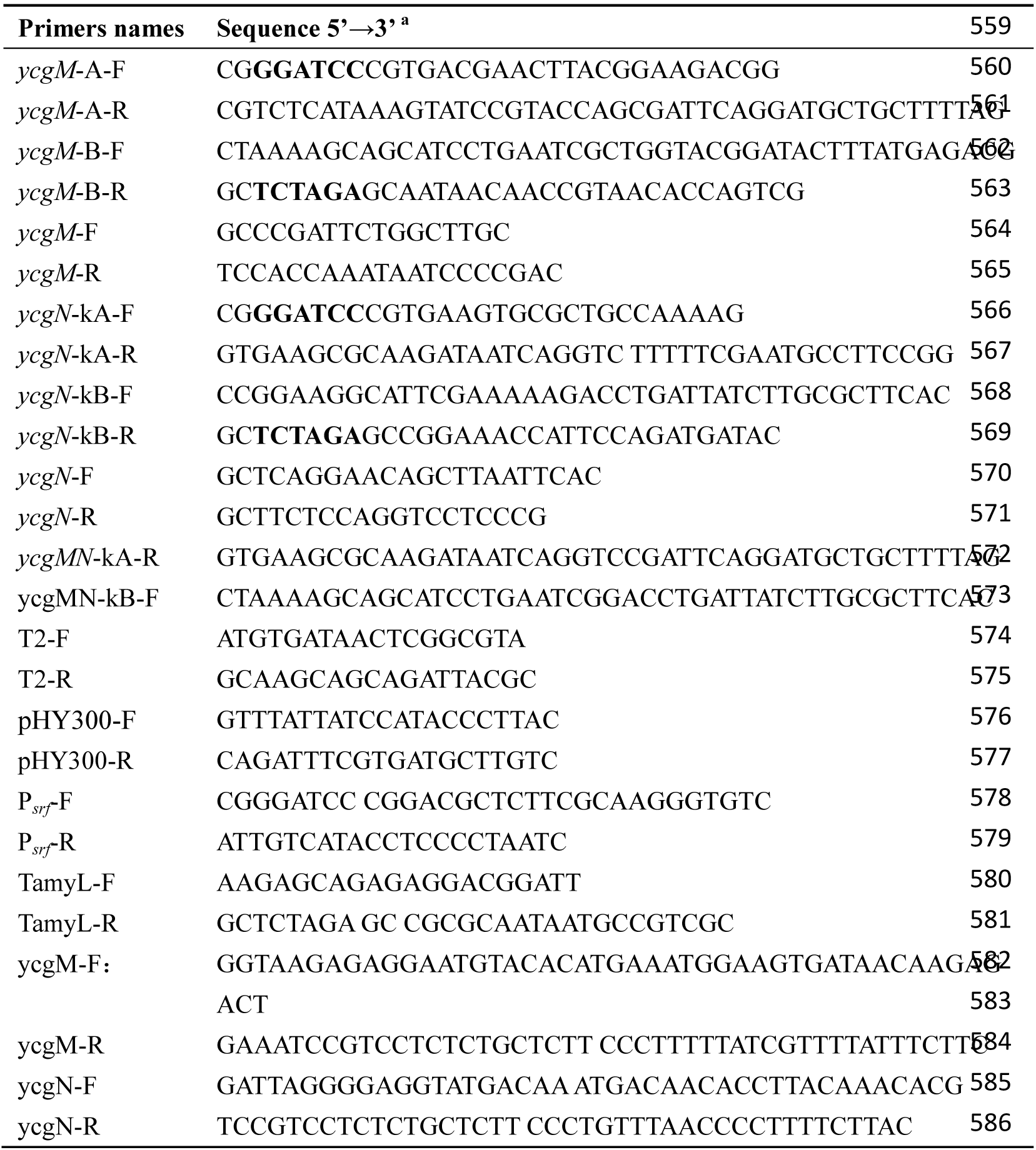
The primers used in this research.

Then, the recombinant vector T2-*ycgN* was transformed into *B. licheniformis* WX-02 by high-osmolality electroporation, according to our previously reported method (37). The transformants were selected by kanamycin resistance and verified by PCR with the primers T2-F and T2-R. Then, the positive colony was cultured in LB medium containing kanamycin (20 mg L^-1^) at 45°C for 8 h to obtain the single-crossover recombinants, and the double-crossover recombinants were screened after serial subculture of single-cross recombinants in LB medium at 37°C. The kanamycin sensitive colonies resulting from the double-crossover event were selected, and confirmed by DNA sequencing with the primers Δ*ycgN*-F and Δ*ycgN*-R. The positive mutant strain was designated as WXΔ*ycgN*. The complement of *ycgN* mutation was generated by introducing the gene *ycgN* into WXΔ*ycgN* at the *amyL* locus, and named as WX Δ*ycgN*-N. Similarly, the *ycgM* deletion, *ycgM* and *ycgN* double deletion strains were constructed with the same method, named WXΔ*ycgM* and WXΔ*ycgMN*, respectively.

The gene expression vector was constructed according to our previously reported method (13). Briefly, the P_*srf*_ promoter (938306) of *B. subtilis* 168, gene *ycgN* (16054241) and the *amyL* terminator (3031010) of *B. licheniformis* WX-02 were amplified by the corresponding primers. The amplified fragments were fused by SOE-PCR. The fused fragment was inserted into pHY300PLK at the restriction enzyme sites *Bam*HI/*Xba*I, resulting in the *ycgN* expression vector, named pHY-*ycgN*. The vectors pHY300PLK and pHY-*ycgN* were then transformed into *B. licheniformis* WX-02, and the recombinant strains were named WX/pHY300 and WX/*ycgN*, respectively. The strain over-expressing *ycgM* was constructed using the similar method, and the recombinant strain was named as WX/*ycgM*.

### Analytical methods

The cell biomass was detected by measuring the absorbance at 600 nm. Briefly, the volume of 2 mL culture broth was centrifuged at 13 700 × *g* for 10 min. The cell pellet was washed three times with 0.85% NaCl solution and re-suspended. The optical density at 600 nm (OD_600_) was measured by a spectrophotometer (Bio-Rad, USA) (38). To determine the *γ*-PGA concentration, three-fold volume of ethanol was added into the supernatant, and centrifuged at 9540 × *g* for 10 min, and resolved with distilled water. The *γ*-PGA concentration was measured by the method of HPLC according to our previous research (14). The concentration of residual glucose was detected by a SBA-40C biosensor analyzer (Institute of Biology, Shandong Province Academy of Sciences, P. R. China) according to the manufacturer’s instructions.

### Determination of proline and P5C concentrations

The cells at logarithmic phase was harvested by centrifugation, washed twice with 0.85% NaCl, and extracted overnight in 5 mL 3% (w/v) aqueous 5-sulphosalicylic acid. Precipitated protein and other debris were removed by centrifugation at 15 000 × *g*, 5 min. To determine the proline content, the volume of 2.0 mL cell extract was reacted with 2.0 mL glacial acetic acid and 2.0 mL acid-ninhydrin (2.5 g ninhydrin was dissolved in the mixed solution of 60 mL glacial acetic acid and 40 mL 6 M phosphoric acid) at 100°C for 1h. Samples were then plunged into ice to stop the reaction, and extracted with 5 mL toluene. The absorbance of toluene phase was separated and measured at 520 nm. The proline concentration was calculated according to the standard curve made by proline standard.

To determine the P5C content, the volume of 1 mL cell extract was added with 0.1 mL trichloroacetic acid, and then 0.5 mL 6 mg mL^-1^ *o*-aminobenzaldehyde (2-AB) was added to the mixture, and insulated at 37°C for 1 h. The mixture was centrifuged at 10 000 × *g* for 10 min. The absorbance of the supernatant was measured at 443 nm, in an l-cm light path. The pyrroline-5-carbosylate concentration (c) was calculated according to Lambert-Beer law: A = ɛ*l*c. The millimolar extinction coefficient (ɛ) of the P5C-oaminobenzaldehyde complex is 2.71 mM^-1^ cm^-1^ (18).

### Determination of ROS

The intracellular ROS levels were measured by the Reactive Oxygen species Assay Kit (Nanjing Jiancheng Bioengineering Institute, P.R. China) according to the manufacturer’s instructions. In brief, the cells were collected by centrifugation at 13 700 × *g* for 10 min, and re-suspended with PBS solution and diluted to OD_600_ = 1.0. The volume of 1 mL cell suspension was added with 10 mM DCFH-DA, and incubated at 37°C for 30 min. The cells were then re-suspended in 1 mL PBS solution, and the relative fluorescence was measured at excitation and emission wavelengths of 485 and 525 nm by using a fluorescence spectrophotometer (Shimadzu, Japan). Untreated cells were used as reference, and the relative ROS amounts were showed by fluorescence intensity (9).

### Determination of ATP concentrations

The intracellular ATP concentration was quantified by a ATP assay kit (Beyotime, China) according to the manufacturer’s instructions (2). Briefly, the cells were lysed by the lysis buffer, and then centrifuged at 12 000 × *g* at 4°C for 5 min, the volume of 20 μL supernatant was mixed with 100 μL luciferase reagent, and the luminance was measured by a luminometer. The ATP concentration was calculated according to the standard curve made by ATP standard, and defined as the content of ATP to the cell dry weight. All assays were performed in triplicate.

### Quantitative real-time PCR (qRT-PCR)

The qRT-PCR assay was conducted according to our previous reported method (13, 36). Briefly, the total RNA was extracted by using the Trizol Reagent (Invitrogen, USA), and DNase I enzyme (TaKaRa, Japan) was applied to degrade trace DNA. The first strand of cDNA was amplified from 0.5 μg of total RNA using the RevertAid First Strand cDNA Synthesis Kit (Thermo, USA) with random primers. The real-time PCR was performed with the Maxima^®^ SYBR Green/ROX qPCR Master Mix (Thermo) following the manufacturer’s instructions. The primers used for amplifying the corresponding genes were listed in **Table S1** (seeing in the Supplementary Material), and 16 S rDNA was used as the reference gene to normalize the data. All the experiments were performed in triplicate. The gene expression levels of recombinant strain were compared with those of wild-type strain after normalization to the reference gene.

## Competing interests

The authors declare that they have no competing interests.

## Athour’s contribution

B Li, Z He and S Chen designed the study. B Li carried out the molecular biology studies and construction of engineering strains. B Li and S Hu carried out the fermentation studies. B Li, D Cai, A Zhu and S Chen analyzed the data and wrote the manuscript. All authors read and approved the final manuscript.

## Acknowledgments

This work was supported by the National Program on Key Basic Research Project (973 Program, No. 2015CB150505), the Technical Innovation Special Fund of Hubei Province (2018ACA149) and the Science and Technology Program of Wuhan (20160201010086).

## Supporting Information

All the primers sequences for RT-qPCR were listed in **Table S1 and Table S2**. This information was available free of charge via the Internet: http://aem.asm.org/.

## Figure caption

**Fig. S1** Effects of H_2_O_2_ dose on cell growth and *γ*-PGA production of WX-02.

